# Cytokines control the physical state of immune tissue

**DOI:** 10.1101/2025.10.28.685215

**Authors:** Debraj Ghose, Thomas C. Ferrante, Donald E. Ingber

**Author notes:** Corresponding author: Donald E. Ingber MD, PhD.

## Abstract

Unlike most solid-like tissues, immune tissue is protean and reconfigurable—its component cells can patrol vast territories, find rare cellular partners, and dynamically self-organize into functional structures like germinal centers or tertiary lymphoid structures. The ability of immune cells to collectively behave as an active material that switches physical state is essential for adaptive immunity. Yet the principles governing the control of the physical properties of immune tissues, and how chemical and mechanical forms of control interplay remain obscure. Inspired by minimal immune cell components that recapitulate germinal center (GC) dynamics, we developed an agent-based model that predicted that a tissue’s ability to program its own physical properties governs the efficiency of antibody evolution. This motivated our search for biomolecular forms of control capable of tuning this material property. Using a tractable in vitro model, we found that CD40L-stimulated human B cells self-organize into active, liquid-like condensates that round, fuse, and internally mix. GC cytokines IL-4 and IL-21 tuned this effective temperature, driving phase transitions from cohesive, liquid-like states to dispersed, gas-like morphologies. These findings demonstrate that cytokines control tissue fluidity, suggesting a previously unrecognized biophysical form of immune regulation.

**Significance Statement:** The body’s immune system is uniquely flexible, allowing cells to organize into temporary structures, like germinal centers or tertiary lymphoid structures, to fight intruders. How this physical flexibility is controlled is a major question. We discovered that the physical state of immune tissue is chemically programmable. We show that self-organized multicellular B cell structures behave like liquid droplets, and we used this framework to demonstrate that signals (cytokines) can modulate their physical properties. This finding identifies a new biophysical axis of immune control, suggesting the body actively tunes its tissue fluidity to optimize the immune response.

## Introduction

Biological tissues are active living materials, whose physical states affect emergent function. The physical principles of liquid-like systems influence function at multiple biological scales, from the dynamics of intracellular biomolecular condensates that organize the cell’s interior (1) to the self-organization of cells into tissues that drive morphogenesis in the embryo (2, 3). However, while most developing tissues transition from liquid-like to solid-like states as they mature (4), immune tissue is a notable exception: it must remain perpetually fluid and protean to carry out its functions of surveillance and response.

This lifelong fluidity is exemplified in germinal centers (GCs), which can form transiently in native secondary lymphoid organs, such as lymph nodes (5), or ectopic tertiary lymphoid structures that can form at cancer or chronic inflammation sites (6–10). In GCs, B cells undergo Darwinian selection to enrich for a subset of B cells that produce high-affinity antibodies (11, 12). Live imaging has shown GCs to be highly dynamic tissue structures that undergo constant cellular rearrangements (13, 14). This is likely because the efficiency of antibody selection depends on how quickly B cells can find antigens, compete for them, locate helper T cells, and shuttle between light and dark zones—all processes fundamentally limited by cellular reorganization dynamics and tissue material properties. The regulation of these dynamics is complex and not fully understood. While B cells can be steered when migrating by chemokine gradients (15) and “decelerate” during direct physical tethering to T follicular helper (TFH) cells (16), it remains unknown if chemical cues could also tune the intrinsic way in which B cells interact with their neighbors in three dimensions. Such a mechanism would represent a distinct and higher-level mode of control, programming the material properties of the tissue by altering multicellular interaction dynamics.

To address the possible existence of this type of physical mechanism of controlling immune cell self-assembly, we first developed a computational model to test the theoretical importance of such regulation within the context of the GC. We then established a simplified experimental system to provide a direct, proof-of-principle test for whether key immune signals can, in fact, reprogram the physical properties of immune tissue. Our findings reveal a previously unrecognized layer of biophysical control in the immune system, where the same signals that dictate immune cell fate also program the active material state of the multicellular collective.

## Results

### Tissue fluidity as a control parameter for B cell evolution

The GC is a transient yet highly organized microanatomical niche within lymphoid organs that functions as an evolutionary machine for B cells (5, 17–20). Within these structures, B cells undergo iterative cycles of mutation, proliferation, and affinity-based selection through dynamic interactions with T follicular helper (TFH) cells and follicular dendritic cells (FDCs) that present antigen (**Fig. 1A**) (12, 13, 17, 20–23). A rich body of theoretical and computational work has provided invaluable insights into the cellular logic and chemokine-driven dynamics of the GC reaction (24–28). Building on this foundation, we sought to first establish the theoretical importance of the tissue’s collective physical properties—a factor distinct from individual cell behaviors. To test the potential link between the physical state of the GC environment and the efficiency of B cell evolution, we developed a minimal spatiotemporal agent-based model that captures the essential features of a GC multicellular reaction.

**Figure 1.**
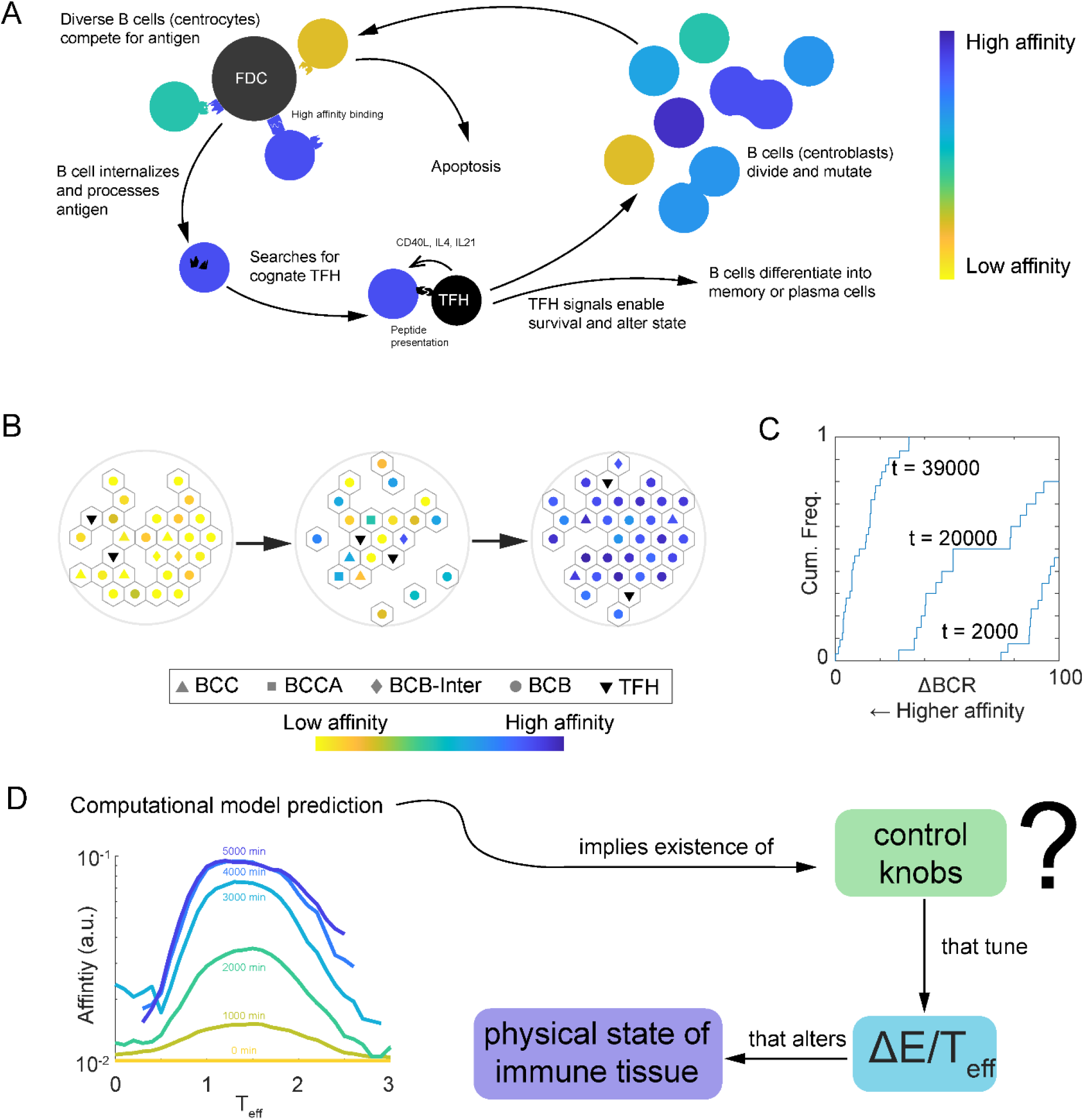
A computational model predicts that B cell evolution is sensitive to tissue fluidity. (A) Schematic of the germinal center (GC) reaction, where B cells undergo iterative cycles of mutation and affinity-based selection through interactions with follicular dendritic cells (FDCs) and T follicular helper (TFH) cells. (B) Snapshots from an agent-based model simulating the GC, showing the progressive enrichment of high-affinity B cells (blue) over lower-affinity B cells (yellowmgreen) over time. (C) The model recapitulates Darwinian selection, as shown by the cumulative frequency plot where the B-cell population shifts toward higher receptor affinity over simulated time. (D) The model predicts that the efficiency of affinity maturation is dependent on the system’s effective temperature (*T*_*eff*_), a parameter controlling tissue fluidity. Affinity maturation is optimal within a specific range, suggesting that the physical state of immune tissue may be a tunable “control knob” for regulating adaptive immunity.

Our computational model populates a hexagonal lattice with B cells in various differentiation states and TFH cells, each simulated as individual agents. The model implements the core GC cycle: B cells in the light zone compete for limited antigen based on their receptor affinity—with higher-affinity cells having a competitive advantage (18, 29, 30). Successfully acquiring antigen and receiving help from TFH cells triggers B cells to migrate to another region of the GC (dark zone), where they proliferate and mutate their receptors before returning to the light zone for another round of selection. This iterative process progressively enriches for high-affinity clones.

In our computational model of the GC, the behavior of each agent is governed by two coupled rule sets. Physical rules dictate tissue rearrangement through Kawasaki Ising dynamics (31–33), where adjacent agents attempt to swap positions with probability determined by the change in system energy. Specifically, the probability of accepting a swap is given by 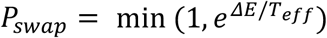 (swaps that decrease energy are always accepted). The change in system energy incorporates multiple contributions: Δ*E* = Δ*E*_*baseline*_ + Δ*E*_*chemotaxis*_ + Δ*E*_*adhesion*_, where baseline energy represents the cost of movement, chemotactic energy drives zone-specific migration toward CXCL13 (light zone) or CXCL12 (dark zone) sources, and adhesion energy promotes cell-cell cohesion. Biological rules control the GC reaction through probabilistic events: antigen acquisition occurs competitively with probability proportional to local CXCL13 concentration, state transitions between B cell subtypes occur at defined rates, cells divide with mutation in regions enriched for CXCL12, and cells undergo apoptosis stochastically.

The overall material state of this simulated tissue emerges from the interplay of these physical and biological processes. While we hold biological rates and interaction strengths constant, we can tune the system’s effective temperature (*T*_*eff*_)—the parameter in the Metropolis acceptance criterion that scales the energetic penalties for movement and thus controls the tissue’s collective fluidity.

We first confirmed that the model recapitulates fundamental Darwinian selection. Over simulated time, the B cell population progressively evolves toward higher affinity as low-affinity clones are outcompeted and eliminated (**Fig. 1B, C**). While not designed for precise quantitative predictions, the model serves as a powerful tool for testing qualitative hypotheses about GC dynamics.

We then used the model to test our central hypothesis: does the physical state of the tissue impact selection efficiency? Running multiple simulations while systematically varying the effective temperature revealed that, at very low temperatures when cells are less motile, affinity maturation proceeds slowly and inefficiently. As temperature increases, selection efficiency improves dramatically, reaching an optimal regime before declining again at very high temperatures, where the system becomes too disordered (**Fig. 1D**).

This theoretical finding, that B cell evolution is sensitive to the system’s effective temperature, is significant. In biological systems refined by natural selection, such parameters are rarely left unregulated. Our model’s prediction thus identifies tissue fluidity as a potential control knob for GC reactions, motivating our subsequent search for biological mechanisms capable of modulating the material state of immune cell collectives.

### B cells stimulated by CD40L form multicellular condensates

Our model’s prediction that B cell evolution is sensitive to effective temperature identifies it as a hypothetical control parameter in the GC multicellular organization reaction. However, the model does not prove that such regulation exists, only that it would be functionally important if the immune system evolved a way to control it. Therefore, the following experiments were designed specifically to test the hypothesis that such biological control knobs exist, by isolating the direct effect of cytokines on the material state of a human immune cell collective.

Addressing this problem within a live GC is currently intractable. In that environment, the tight coupling of chemical signaling with physical cell-cell contact and the simultaneous triggering of complex cell fate programs (e.g., proliferation and selection) make it challenging to isolate the direct effect of any single cytokine on the material state of the entire tissue. Instead, we required a simplified system where we could directly measure the material state of an immune cell collective in response to specific chemical cues. We therefore leveraged a well-described phenomenon where stimulating CD40 receptors in primary human B cells, which mimics the initial TFH engagement, causes them to spontaneously self-organize into multicellular aggregates in suspension culture (**Fig. 2A**) (34–36). While these aggregates are known to form, their physical properties are poorly understood.

**Figure 2.**
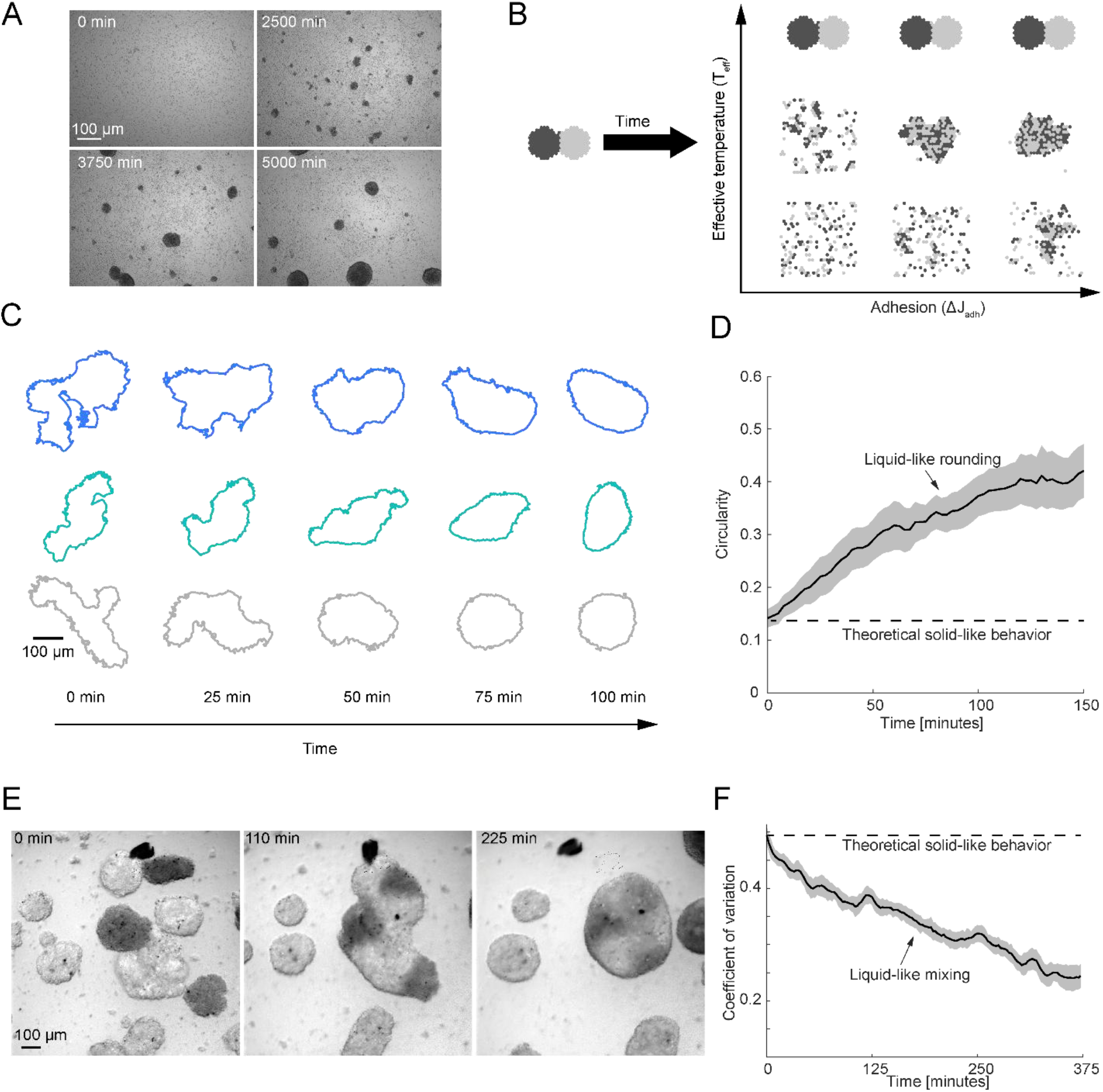
CD40L-stimulated B cells self-organize into liquid-like multicellular condensates. (A) A representative brightfield image of multicellular aggregates formed by primary human B cells stimulated with CD40L. (B) A computational simulation shows two distinct populations of cells (dark gray and light gray) spontaneously intermixing over time, a characteristic of a fluid system. (C) Time-lapse outlines of three representative B-cell aggregates show that initially irregular structures round up over 100 minutes, indicating the presence of an effective surface tension that minimizes surface area. (D) Quantification of rounding behavior. A plot of circularity versus time shows a steady increase, consistent with liquid-like relaxation dynamics. n = 11 and standard error of the mean is shown gray. (E) Time-lapse microscopy reveals that separate B-cell aggregates fuse upon contact and their constituent cells intermix, further demonstrating their liquid-like nature. (F) Quantification of the intermixing process shows a decrease in the coefficient of variation over time, confirming the internal fluidity of the B-cell condensates. n = 17 and standard error of the mean is shown gray.

To adopt this simplified in vitro model system, we first set out to characterize its physical nature within the framework of soft matter physics. Given that germinal centers are known to be highly dynamic sites of constant cellular rearrangement, we hypothesized that these self-organized homotypic B cell aggregates may also rest within a similar liquid-like physical state. To explore this possibility theoretically, we used a simplified computational model that considers only a single cell type and homotypic adhesion, representing the aggregates formed by B cells. This model predicts a phase diagram where the collective state of the cells is governed by the interplay between cell-cell adhesion energy (Δ*J*_*adhesion*_) and an effective temperature (*T*_*eff*_) that controls cell motility (**Fig. 2B**). At the extremes, low temperatures and high adhesion promote solid-like crystalline states, while high temperatures and low adhesion lead to dispersed, gas-like states. Between these regimes, the model predicts the existence of liquid-like condensates, which are characterized by their ability to exhibit rounding behavior and internal cellular rearrangements.

To experimentally characterize multicellular B cell structures in the context of the phase diagram in **Fig. 2B**, we used multimerized CD40L (6.25 µ gmmL; 75 UmmL) to induce B cell aggregate formation (37, 38) and imaged their dynamics over time using live-cell microscopy (**Movie S1**). Tracking the boundaries of these structures revealed that irregularly shaped structures rounded over time, a hallmark of surface-tension-driven rounding observed in liquid droplets (**Fig. 2C**). This rounding behavior was quantified by a steady increase in circularity over time, approaching a fluid equilibrium state rather than a jammed solid one (**Fig. 2D**), suggesting that multicellular B cell structures behave like liquid droplets.

Another defining characteristic of a liquid is internal fluidity, which allows constituent components to mix and rearrange. To test this, we prepared two populations of B cells labeled with distinct fluorescent dyes. When co-cultured, aggregates of different colors readily fused upon contact, and their constituent cells thoroughly intermixed over several hours, confirming their liquid-like fluidity (**Fig. 2E** and **Movie S2**). This internal mixing was confirmed by a decrease in the coefficient of variation over time, again consistent with a liquid-like system (**Fig. 2F**).

Taken together, direct experimental evidence of both effective surface tension and internal fluidity establishes that stimulated B cells self-organize into active liquid-like droplets. We term these structures “B-Lymphocyte Organized Multicellular Blobs” (BLOMBs) for simplicity of description. This characterization provided us with the ideal platform to test our central hypothesis that TFH cytokines which are known to alter B cell state in the GC may also tune the material properties of immune tissue.

### TFH-derived cytokines tune the physical state of BLOMBs

Having established BLOMBs as multicellular condensates, we next sought to identify the biological control knobs capable of tuning their physical state. Our computational model predicted that decreasing Δ*E*/*T*_*eff*_ by increasing *T*_*eff*_ would decrease the circularity of multicellular condensates (**Fig. 3A**). This provided a clear visual signature to look for in our experiments.

**Figure 3.**
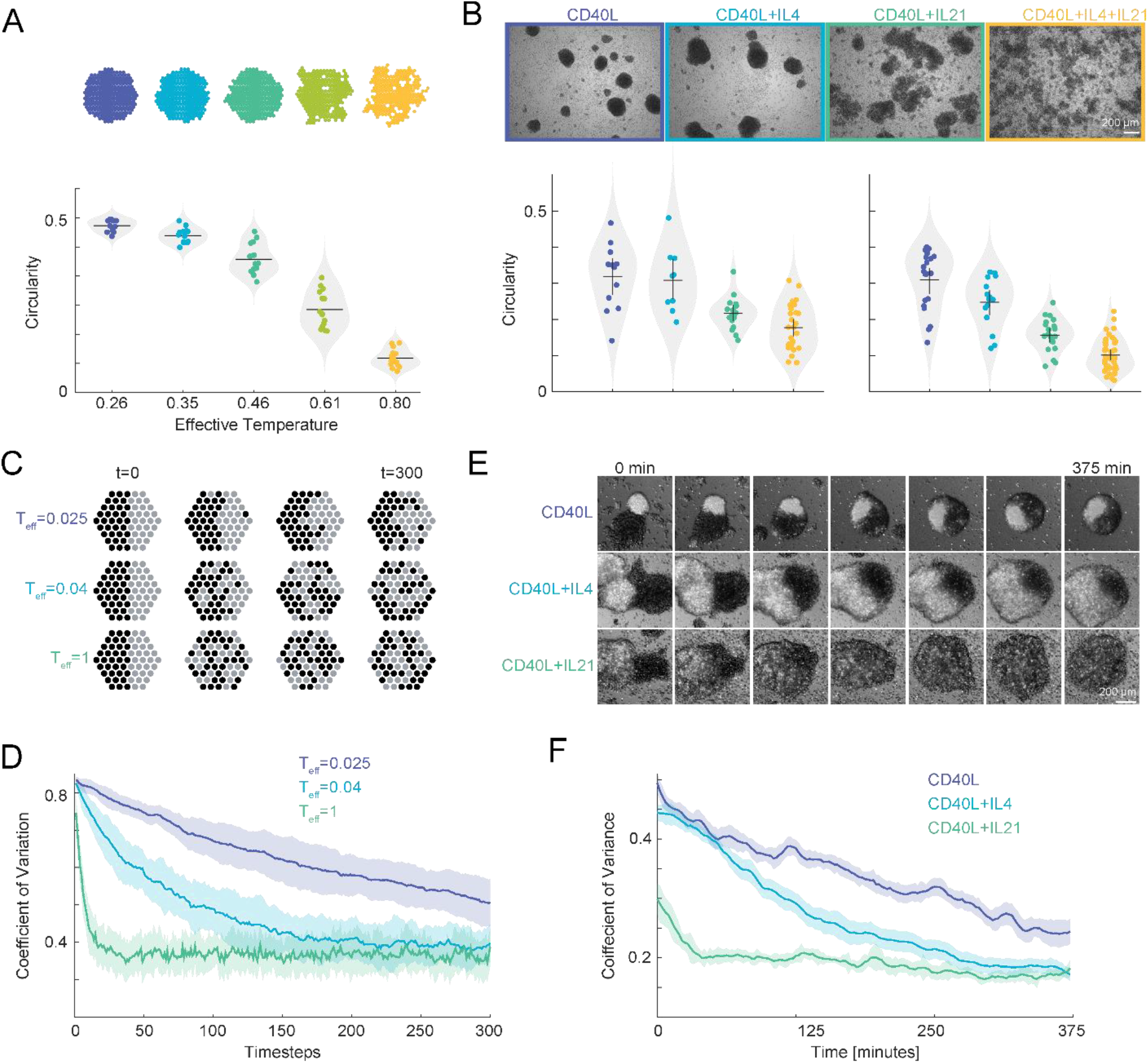
TFH-derived cytokines tune the physical state of B-cell collectives. (A) Agent-based Kawasaki-Ising model that only considers one cell type and adhesion. The model predicts that increasing T_eff_, which lowers ΔE/T_eff_, leads to a loss of cohesion, causing simulated cell aggregates to become more dispersed and less circular. (B) Experimental validation shows that TFH cytokines modulate the physical state of B-cell aggregates across two different human donors. While aggregates treated with CD40L alone or with IL-4 remain cohesive, the addition of IL-21 induces a transition to a dispersed, gas-like state, characterized by a significant decrease in circularity. This phenocopies the effect of increasing effective temperature in the model. The horizontal black lines are the bootstrapped means, and the black vertical bars show the 95% bootstrap confidence interval of the mean (nboot = 1000, resampling particles within each condition). (C) Snapshots from the agent-based model at different effective temperatures ( T_eff_ ) illustrate the transition from a condensed to a dispersed state over time. (D) Quantification from the simulation shows that the coefficient of variation, a measure of intermixing, decreases more rapidly at higher effective temperatures. (E) Experimental time-lapse microscopy of B-cell aggregates shows their structural evolution over 375 minutes under different cytokine conditions. CD40L n=17, CD40L+IL4 n=17, CD40L+IL4+IL21=21, and shaded regions are 95% confidence intervals of mean. (F) Experimental quantification confirms the model’s prediction: the coefficient of variation decreases most rapidly in the presence of IL-21, indicating that this cytokine increases the system’s effective temperature and drives the collective toward a more fluid state.

Thus, we sought to identify biological factors that could induce this predicted physical change— namely, a transition to a more dispersed, less circular state. Our BLOMB system is formed by stimulating B cells with CD40L, which mimics the survival signal B cells receive from TFH cells (**Fig. 1A**). We reasoned that the other key signals delivered by TFH cells during this interaction might serve as the ‘control knobs’ capable of tuning the collective’s physical state. We therefore focused on IL-4 and IL-21, two canonical TFH-derived cytokines that are known to act in concert with CD40L to direct B cell fate, driving proliferation, survival, and differentiation (39–44). The signaling pathways downstream of the IL-4 and IL-21 receptors are known to intersect with pathways that regulate the actin cytoskeleton, contractility, and cell adhesion—the very components that govern a cell’s mechanical properties and biophysical interactions with other cells (45–48). We therefore hypothesized that these potent, fate-directing signals might also function to modulate the collective physical state.

To test this hypothesis, we perturbed BLOMBs from two donors with IL-4 (2.5 µgmmL) and IL-21 (1.25 µgmmL). These saturating concentrations were chosen to ensure maximal signaling and provide a clear proof-of-principle test for whether these cytokines could modulate the collective’s physical state. Imaging revealed striking morphological changes: while BLOMBs treated with CD40L alone or with IL-4 remained cohesive, the addition of IL-21 induced a transition to a dispersed, gas-like state. This was quantified by a significant decrease in aggregate circularity, a phenotype that was amplified when both cytokines were present (**Fig. 3B**). This cytokine-driven shift from a cohesive ‘liquid-like’ to a dispersed ‘gas-like’ state phenocopies the effect of decreasing Δ*E*/*T*_*eff*_ in our model (**Fig. 3A**).

To further investigate the system dynamics, we compared time-lapse data from both simulations and experiments. Our model shows that increasing the effective temperature not only changes the final state but also accelerates the dynamics towards a more dispersed configuration, as seen in simulation snapshots (**Fig. 3C**) and quantified by a more rapid decrease in the coefficient of variation (**Fig. 3D**). This theoretical prediction was mirrored in our experiments. Time-lapse microscopy revealed the structural evolution of the BLOMBs under different cytokine conditions (**Fig. 3E**). Quantification confirmed that the coefficient of variation decreased most rapidly and reached a lower plateau in the presence of IL-21, indicating that this cytokine increases the system’s effective temperature and drives the collective toward a more fluid, gas-like state (**Fig. 3F**).

These results demonstrate that TFH-derived cytokines can modulate the physical state of B-cell collectives. By tuning Δ*E*/*T*_*eff*_, these signals fluidize the tissue, directly linking the chemical signaling environment to the material properties our model identified as crucial for the GC reaction.

## Discussion

Our findings reveal a biophysical layer of control in immune tissue regulation where the same cytokine signals that determine B cell fate and promote Darwinian selection in the germinal center (GC) also control the physical state of cell collectives. We established this principle by combining computational modeling with live cell imaging. First, our *in silico* model, which simulates the essential features of the GC, predicted that the efficiency of B cell evolution is optimally tuned by the system’s ‘effective temperature’. Second, to determine if a biological mechanism for tuning this physical parameter exists, we experimentally perturbed a reductionist *in vitro* system of B-cell condensates, which we term B-Lymphocyte Organized Multicellular Blobs (BLOMBs). This in vitro experimental system is not intended to replicate the complex architecture of an in vivo GC. Rather, its value lies in its simplicity, which enabled a direct, proof-of-principle demonstration that the TFH cytokines IL-4 and IL-21 can reprogram the material properties of a B cell collective, tuning its state from cohesive to dispersed form. Thus, our work establishes the existence of a new form of biophysical control in immune tissues (**Fig. 4**). How the immune system precisely leverages this tunability to optimize functions like affinity maturation in vivo remains a key question for future investigation.

**Figure 4.**
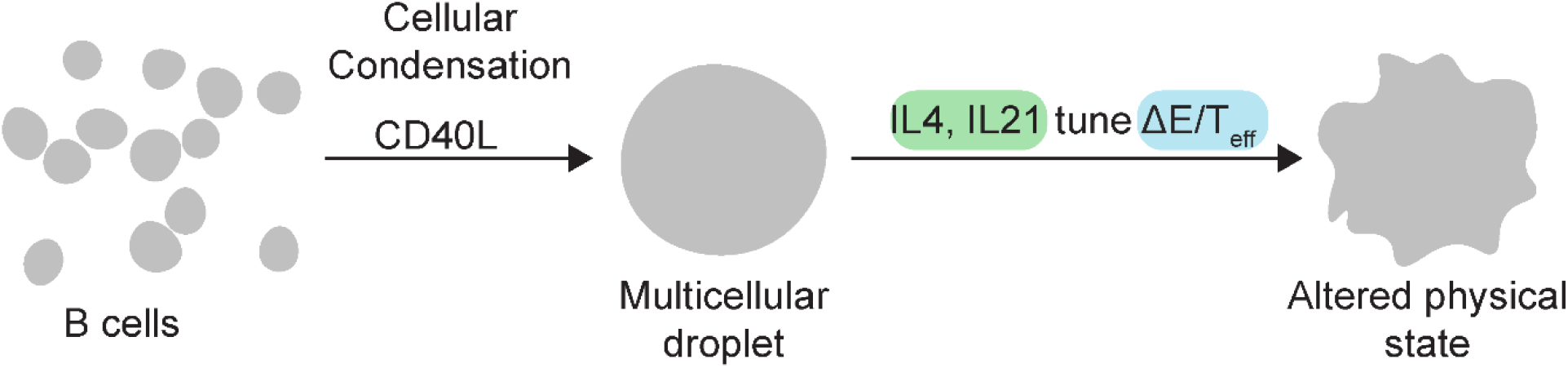
A model for cytokine control of immune tissue physical state. This schematic summarizes the paper’s central findings. Initially dispersed B cells, upon stimulation with CD40L, undergo a form of cellular condensation to self-organize into cohesive, liquid-like multicellular droplets. These droplets represent a baseline physical state. The introduction of T follicular helper (TFH) cell-derived cytokines, such as IL-4 and IL-21, acts as biological control knobs, tuning Δ*E*/T_eff_. This cytokine signaling can drive a phase transition, altering the collective’s material properties from a cohesive liquid-like state to a more dispersed, gas-like state, thereby providing a mechanism for physically programming immune tissue.

This work builds on the growing understanding that tissues can behave as active materials with liquid-like properties, a concept well-established in the study of embryonic development (2). The key distinction, however, is that while embryonic tissues typically solidify upon maturation (4), our findings demonstrate that immune tissue not only maintains a fluid state throughout life but that this state is actively and chemically programmable. This concept of a tunable material state draws a powerful parallel to the regulation of intracellular biomolecular condensates (1). Although the underlying molecular drivers and physical scales are distinct, the analogy is instructive: in both systems, the collective interactions of individual components give rise to a condensed phase whose material properties are tuned to regulate a specific biological function.

We therefore propose that immune cell assemblies are a form of programmable active matter, distinct from other tissues that typically solidify during development. Having identified a core principle of this programmability in our simplified system, a key future challenge is to understand how these rules apply within the complex 3D architecture of diverse lymphoid structures *in vivo*. For example, the fluidity of the GC may be actively regulated to enhance the search-and-selection process during affinity maturation, ensuring that B cells can efficiently find antigen and connect with helper T cells.

Beyond the GC, this principle may govern the dynamics of ectopic tertiary lymphoid structures (TLSs) that form at sites of chronic inflammation or cancer (6–10). The ability to tune the local tissue fluidity could impact immune cell infiltration, residency, and anti-tumor activity, as well as lead to the identification of new therapeutic targets. Similarly, this biophysical control may be crucial in shaping mucosal immunity within specialized niches like inducible bronchus-associated lymphoid tissue or IgA-producing intestinal cryptopatches (49, 50). In these environments, controlling the collective state of lymphocytes could be essential for maintaining barrier integrity while allowing for rapid and flexible immune responses. This active matter immunology perspective shifts the focus from purely biochemical signaling to a model where the physical state of the tissue is itself a regulated variable critical for immune function.

The principles uncovered in this study also suggest new avenues for the rational design of immune tissues. The self-organizing and physically tunable nature of BLOMBs provides a foundation for a “bottom-up” approach to tissue engineering, where cytokine inputs can be used to program the collective state and geometry of a multicellular tissue construct. For example, by harnessing this cytokine-mediated control over the material state, it may be possible to engineer synthetic lymphoid organoids *ex vivo*.

Such engineered immune tissues could serve as powerful platforms for fundamental research, allowing for the deconstruction of complex immune processes in a highly controlled environment. They could also accelerate translational work by enabling high-throughput screening of immunomodulatory drugs or serving as systems to produce therapeutic antibodies. Looking forward, this work paves the way for fabricating active biological materials with prescribed and reconfigurable functions, hinting at a future where engineered immune tissues could be designed for therapeutic applications. In short, our work clarifies that immune tissue is a protean, programmable active material, and that tuning its physical state represents a powerful and previously unrecognized axis for immunomodulation.

## Materials and Methods

Primary human B lymphocytes were isolated from PBMCs from donor leukocyte collars via immunomagnetic isolation and cultured as described in SI Appendix. Live-cell time-lapse imaging was performed using brightfield and fluorescence microscopy, and image analysis used custom scripts in MATLAB 2024a, as detailed in SI Appendix. The agent-based and Kawasaki-Ising computational models were implemented in MATLAB 2024a as described in SI Appendix.

## Acknowledgments

We thank Aditya Patil, Pranav Prabhala, Min Wu, Girija Goyal, Yunhao Zhai and members of the Ingber lab for generously sharing reagents and techniques. We are also grateful to Duanne Wesemann and Harikesh Wong for valuable feedback and discussions. This work was supported by the Sloan Matter-to-Life program and the Wyss Institute for Biologically Inspired Engineering at Harvard University.

## Author Contributions

D.G. conceptualized the project, designed the methodology, performed the investigation and formal analysis, and wrote the original draft and subsequent revisions. T.C.F. contributed to the methodology. D.E.I. acquired funding, contributed to conceptual discussions, and participated in writing, review, and editing.

## Competing Interests

The authors declare no competing interests.

## Data Availability Statement

All primary data and analysis code supporting the findings of this study are publicly available on GitHub at https://mmgithub.commDebrajGhosemcytokine_control_blomb. Further information and requests for resources or materials should be directed to the corresponding author (D.E.I.).

## Notes

https://github.com/DebrajGhose/cytokine_control_blomb

### Competing Interest Statement

The authors have declared no competing interest.

### Summary of Updates

Energy term in Figure 4 relabeled from J to E.

