## Supplementary material for "Cytokines control the physical state of immune tissue": SI Appendix

### Supplemental Material for BLOMB paper

xe *[1]

[1]Wyss Institute for Biologically Inspired Engineering, Harvard University, Boston, MA, 02215, USA

March 2025

### 1 Computational Model of Germinal Center Dynamics

We developed a two-dimensional computational model to simulate the dynamics of a germinal center (GC) by representing individual cells on a hexagonal lattice. The model employs a multi-component Kawasaki dynamics algorithm, adapted from the Ising model framework [1–4], to govern cell movement. This framework treats cell movement as a spin-exchange process where cells can swap positions based on energy considerations, integrating both adhesive interactions and chemotactic gradients. The model also has stochastic rules for biological processes such as cell proliferation, differentiation, and selection. The simulation was implemented in MATLAB.

#### 1.1 Spatial Framework and Environment

The simulation domain consists of a two-dimensional hexagonal lattice, with cells occupying discrete lattice sites. The distance between the centers of adjacent sites is $d_{hex} = 10$ µm (twice the cell radius). The model defines three concentric circular boundaries:

- A germinal center radius of $R_{GC} = 30$ µm for chemokine source placement
- A movement boundary at $R_{boundary} = 100$ µm beyond which cells cannot move
- An escape boundary at $R_{escape} = 120$ µm beyond which cells are removed from the simulation

To establish the characteristic light zone (LZ) and dark zone (DZ) polarization, we impose two static, opposing chemokine gradients. The CXCL13 source (for the LZ) is positioned at $(-0.9 \times R_{GC}, 0)$ and the CXCL12 source (for the DZ) at $(0.9 \times R_{GC}, 0)$. The concentration $C$ of each chemokine is a function of the distance $d$ from its source point, following a Hill-like equation:

$$C(d) = \frac{K^n}{K^n + d^n} \tag{1}$$

where $K = 20$ µm is the half-maximal concentration distance and $n = 2$ is the Hill coefficient.

#### 1.2 Cell Types and States

Each site on the lattice can either be empty or occupied by a single cell. Cells are distinguished by their type, $\tau$, which determines their behavior. The model includes five types:

- **Centroblast (BCB):** Proliferating B cells that are attracted to CXCL12 in the dark zone. These cells have a division limit to prevent unlimited expansion.
- **Intermediate Centroblast (BCB-Inter):** A transient state for B cells re-entering the cell cycle after receiving T-cell help. Upon transition to this state, the cell's division counter is reset.
- **Centrocyte (BCC):** Non-dividing B cells attracted to CXCL13 in the light zone to seek antigen.
- **Antigen-Captured Centrocyte (BCCA):** Centrocytes that have acquired antigen and now seek T-cell help.
- **T Follicular Helper (Tfh) Cell:** Mobile cells in the light zone that provide survival signals.

#### 1.3 Kawasaki Dynamics and Cell Movement

Cellular migration and sorting are emergent properties governed by Kawasaki (spin-exchange) dynamics. The system's total energy is given by a Hamiltonian that depends on pairwise interactions between neighboring sites:

$$H = \sum_{\langle i,j \rangle} J_{\sigma_i,\sigma_j} \tag{2}$$

where the sum runs over all nearest-neighbor pairs $\langle i, j \rangle$, and $\sigma_i$ represents the occupant at site $i$.

At each simulation time step ($\Delta t = 1$ min), every cell is given one opportunity to move in a randomized order. A cell at site $i$ randomly selects one of its six hexagonal neighbors as a potential target site $j$. The system calculates the energy change $\Delta E$ for swapping the occupants of sites $i$ and $j$ (which could be another cell or empty space).

The total energy change includes contributions from cell-cell adhesion, chemotaxis, and a baseline movement cost:

$$\Delta E = \Delta E_{adhesion} + \Delta E_{chemotaxis} + J_{baseline} \tag{3}$$

The adhesion energy change is calculated as:

$$\Delta E_{adhesion} = J_{adhesion} \cdot (\eta_{after} - \eta_{before}) \tag{4}$$

where $J_{adhesion} = -1.0$ is the adhesion energy (negative for attraction) and $\eta$ represents the number of neighboring cells.

The chemotaxis energy for a cell of type $\tau$ at position $\vec{x}$ is:

$$E_{chemotaxis}(\vec{x}, \tau) = -\lambda_{chemo} \cdot C_{attractant}(\vec{x}) \tag{5}$$

where $\lambda_{chemo} = 5.0$ is the chemotaxis strength and $C_{attractant}$ is the concentration of the relevant chemokine (CXCL13 for BCC, BCCA, and Tfh cells; CXCL12 for BCB and BCB-Inter cells).

The baseline movement cost $J_{baseline} = 1.0$ represents a general energetic penalty for any movement.

The proposed swap is accepted with probability:

$$P(\Delta E) = \min\left(1, e^{-\Delta E / T_{eff}}\right) \tag{6}$$

where $T_{eff}$ is the effective temperature. For a swap between two cells, $T_{eff}$ is the average of their individual motility temperatures. For a cell moving to an empty site, it equals the cell's temperature. In the Kawasaki framework, this effective temperature controls the balance between energy-driven organization (at low $T$) and random thermal motion (at high $T$), determining whether the system exhibits solid-like, liquid-like, or gas-like behavior.

#### 1.4 Biological Processes and State Transitions

In addition to movement, cells undergo state changes based on stochastic rules that model key biological events:

- **Antigen Acquisition:** A BCC in the light zone becomes a BCCA with probability $P_{acq} = k_{antigen} \cdot C_{CXCL13} \cdot \Delta t$, where $k_{antigen} = 0.005$. The model preferentially targets neighboring BCC cells with the lowest BCR affinity within a 1.5 hex-spacing radius.
- **T-Cell Help:** A BCCA directly adjacent to a Tfh cell transitions to the BCB-Inter state and resets its division counter to 0.
- **Differentiation:** Cells transition between states with fixed rates:
  - BCB-Inter → BCB with rate $k_{BCB\_inter\_to\_BCB} = 1/600 \text{ min}^{-1}$
  - BCB → BCC with rate $k_{BCB\_to\_BCC} = 1/300 \text{ min}^{-1}$
- **Proliferation and Mutation:** BCB cells divide with probability $P_{divide} = k_{divide} \cdot C_{CXCL12} \cdot \Delta t$ (where $k_{divide} = 1/240 \text{ min}^{-1}$), provided:
  - An empty adjacent lattice site is available
  - The cell has not exceeded its division limit (6 divisions)
  - The total population is below the carrying capacity (50 cells)

  Upon division, both parent and daughter cells undergo somatic hypermutation with probability $p_{mutation} = 0.75$, modifying their BCR affinity by a value drawn from $\mathcal{N}(0, \sigma_{mut}^2)$ where $\sigma_{mut} = 8.0$.
- **Apoptosis:** Unselected BCC and BCCA cells die with rate $k_{death} = 1/600 \text{ min}^{-1}$.

#### 1.5 Population Control Mechanisms

The model implements two key mechanisms to control population dynamics. First, each BCB cell can divide a maximum of 6 times. This counter is inherited by daughter cells but resets when a cell receives T-cell help. Second, the total population is limited to 50 cells.

##### 1.6 Simulation Initialization

Simulations begin with $N_{B\_initial} = 20$ centrocytes (BCC) and $N_{Tfh} = 2$ Tfh cells. Tfh cells are placed near the CXCL13 source (light zone center) on the hexagonal grid, while B cells are randomly distributed across valid hexagonal lattice positions within the movement boundary. Each simulation runs for $T_{total} = 50,001$ minutes with data logged at 10,000-minute intervals.

Table 1: Simulation Parameters for Germinal Center Model

| Parameter | Value | Description |
|---|---|---|
| | | **Spatial Parameters** |
| $R_{GC}$ | 30 µm | Germinal center radius (for chemokine sources) |
| $R_{boundary}$ | 100 µm | Movement boundary radius |
| $R_{escape}$ | 120 µm | Escape boundary (cell removal) |
| $r_{cell}$ | 5 µm | Approximate radius of a single cell |
| $d_{hex}$ | 10 µm | Spacing between hexagonal lattice sites |
| | | **Temporal Parameters** |
| $\Delta t$ | 1 min | Duration of a single time step |
| $T_{total}$ | 50,001 min | Total simulation time |
| | | **Energy and Motility Parameters** |
| $J_{adhesion}$ | -1.0 | Adhesion energy between cells |
| $J_{baseline}$ | 1.0 | Baseline movement energy cost |
| $\lambda_{chemo}$ | 5.0 | Strength of chemotactic energy |
| $T_{BCC}$ | 1.0 | Motility temperature of Centrocytes |
| $T_{BCCA}$ | 1.0 | Motility temperature of Ag-Captured Centrocytes |
| $T_{BCB\_inter}$ | 1.0 | Motility temperature of Intermediate Centroblasts |
| $T_{BCB}$ | 1.0 | Motility temperature of Centroblasts |
| $T_{Tfh}$ | 1.0 | Motility temperature of Tfh cells |
| | | **Chemokine Gradient Parameters** |
| $K$ | 20 µm | Half-maximal concentration distance |
| $n$ | 2 | Hill coefficient for gradient steepness |
| | | **Biological Event Rates** |
| $k_{divide}$ | 1/240 $\text{min}^{-1}$ | Base rate of cell division |
| $k_{death}$ | 1/600 $\text{min}^{-1}$ | Rate of apoptosis |
| $k_{BCB\_to\_BCC}$ | 1/300 $\text{min}^{-1}$ | Transition rate from BCB to BCC |
| $k_{BCB\_inter\_to\_BCB}$ | 1/600 $\text{min}^{-1}$ | Transition rate from BCB-Inter to BCB |
| $k_{antigen}$ | 0.005 | Base rate of antigen acquisition |
| | | **Mutation Parameters** |
| $p_{mutation}$ | 0.75 | Probability of mutation upon division |
| $\sigma_{mut}$ | 8.0 | Standard deviation of BCR score change |
| | | **Population Control** |
| Division limit | 6 | Maximum divisions per BCB lineage |
| Carrying capacity | 50 | Maximum total cell population |
| | | **Initial Conditions** |
| $N_{B\_initial}$ | 20 | Initial number of B cells |
| $N_{Tfh}$ | 2 | Initial number of Tfh cells |

#### 2 Phase Diagram Simulations

To understand the fundamental physics underlying our Kawasaki dynamics model, we performed simplified simulations exploring the phase behavior of cellular aggregates. These simulations examined how varying adhesion strength ($J$) and effective temperature ($T_{eff}$) affects the collective behavior of cells, revealing transitions between gas-like (dispersed), liquid-like (cohesive but mobile), and solid-like (rigid) phases.

#### 2.1 Two-Blob Interaction Model

We simulated the interaction between two initially separated circular blobs of cells to map the phase diagram of the system. Each blob had radius $R_{blob} = 50$ µm and contained approximately 75-100 cells arranged on the hexagonal lattice. The blobs were initially separated by a distance of $2R_{blob}$ with centers at $(-R_{blob}, 0)$ and $(R_{blob}, 0)$.

The simulation used simplified energy parameters:

- Universal adhesion energy $J$ between any two cells (varied from -0.1 to -20)
- Baseline movement cost $J_{baseline} = 0.05$
- No chemotaxis or other directional forces

Cells could either move to empty neighboring sites or swap positions with adjacent cells, following the standard Kawasaki dynamics described in Section 1. The system was confined to a radius of $3.5R_{blob}$ to prevent unlimited dispersal. Simulations ran for 5,000 time steps, sufficient to reach steady-state configurations.

#### 2.2 Phase Behavior

The simulations revealed three distinct phases as a function of adhesion strength and temperature:

**Gas Phase** (High $T_{eff}$, Weak $|J|$): Cells disperse throughout the available space with minimal clustering. The thermal energy exceeds the adhesive interactions, preventing stable aggregate formation. In this regime, the two initial blobs dissolve and cells explore the entire accessible area randomly.

**Liquid Phase** (Intermediate $T_{eff}$ and $|J|$): Cells maintain cohesive clusters while retaining internal mobility. The two blobs can merge if they come into contact through random fluctuations, forming a single larger aggregate. Within aggregates, cells continuously rearrange positions, allowing for shape changes and internal mixing. This phase is most relevant for biological systems where both cohesion and plasticity are required.

**Solid Phase** (Low $T_{eff}$, Strong $|J|$): Cells form rigid structures with minimal internal rearrangement. The two blobs remain separate even when in contact, as the energy barrier for mixing exceeds available thermal fluctuations. Individual cells become locked in place by their neighbors, preventing both shape changes and mixing.

### 3 Metrics for Characterizing Cellular Dynamics

To quantitatively assess the behavior of cellular aggregates in our simulations, we developed two complementary metrics: circularity for measuring shape evolution and a mixing coefficient for quantifying cell type segregation.

#### 3.1 Circularity Metric

Circularity quantifies how closely an aggregate's shape resembles a perfect circle, providing insight into surface tension-driven shape relaxation. We define circularity as:

$$\text{Circularity} = \frac{4\pi A}{P^2} \tag{7}$$

where $A$ is the area of the aggregate and $P$ is its perimeter. This metric ranges from 0 to 1, with 1 representing a perfect circle.

##### 3.1.1 Implementation Details

For cells on a hexagonal lattice, the implementation requires specific methods for area and perimeter calculation. The total **area** $(A)$ is found by multiplying the number of cells $(N)$ by the area of a single hexagon, given by $A = N \cdot \frac{3\sqrt{3}}{2}(r_{hex})^2$, where $r_{hex} = d_{hex}/2$. To find the **perimeter** $(P)$, we first identify boundary cells, which are defined as those with fewer than 6 neighbors. This boundary detection is achieved by checking if any of a cell's 6 hexagonal neighbor positions are unoccupied, using distance-based checking with a tolerance $< 10^{-6}$ to handle floating-point comparisons robustly. The total perimeter is then computed by summing the lengths of the exposed edges of all identified boundary cells, where each exposed edge has a length of $d_{hex}/\sqrt{3}$.

###### 3.1.2 Circularity Evolution Simulations

To demonstrate the utility of this metric, we simulated the evolution of initially irregular blobs containing approximately 50 cells. The irregular shapes were generated using a biased growth algorithm that created connected but non-circular aggregates with a controlled irregularity parameter (0 = circle, 2 = highly irregular).

##### 3.2 Mixing Coefficient

To quantify the degree of mixing between two cell types within an aggregate, we developed a coefficient of variation (CV) based on local composition averaging.

###### 3.2.1 Local Averaging Method

For each cell, we calculate the local fraction of type-1 cells considering both the cell itself and its immediate hexagonal neighbors:

$$f_i^{local} = \frac{n_{type1}^{(i)}}{n_{total}^{(i)}} \tag{8}$$

where $n_{type1}^{(i)}$ is the number of type-1 cells (including self) and $n_{total}^{(i)}$ is the total number of cells in the local neighborhood of cell $i$.

The mixing coefficient is then computed as:

$$CV = \frac{\sigma(f^{local})}{\mu(f^{local})} \tag{9}$$

where $\sigma$ and $\mu$ are the standard deviation and mean of local fractions across all cells. This metric captures local heterogeneity: a **CV $\approx$ 1** indicates complete segregation (each cell surrounded only by the same type), a **CV $\approx$ 0** represents perfect mixing (each cell has proportional representation of both types), and an **intermediate CV** signifies partial mixing with some clustering.

###### 3.2.2 Mixing Dynamics Simulations

We simulated mixing within a single circular blob (radius 40 µm) initially divided into left (type 1) and right (type 2) halves. Cells could only swap positions (no moves to empty space) to preserve the aggregate shape while allowing internal rearrangement.

#### 4 Experimental Analysis of BLOMB Circularity and Mixing

To validate our computational predictions, we performed quantitative analysis of BLOMB morphology and cell mixing dynamics using time-lapse microscopy of experimentally generated aggregates. This section describes the image analysis methods used to extract circularity and mixing metrics from microscopy data.

##### 4.1 Cell Culture Media

All primary cells were cultured in RPMI 1640 medium supplemented with 10% fetal bovine serum (FBS) and 1% antibiotic solution. This formulation is referred to as "complete culture media" in the text.

##### 4.2 Isolation of Primary Human Naïve B Cells

Primary human peripheral blood mononuclear cells (PBMCs) were isolated from leukapheresis collars obtained from Brigham and Women's Hospital. The blood product was first diluted with an equal volume of PBS or RPMI medium and then slowly layered onto Lymphoprep (Catalog #07801) density gradient medium. The samples were centrifuged at 300 x g for 25 minutes at room temperature with minimal acceleration and deceleration settings. The resulting buffy coat, containing the PBMC fraction, was carefully collected.

These cells were washed by adding them to 10 mL of media and centrifuging at 120 x g for 10 minutes to remove excess platelets. After aspirating the supernatant, the PBMC pellet was centrifuged again at 300 x g for 5 minutes. The cells were then resuspended at a concentration of 5 x $10^7$ cells/mL in the recommended sorting buffer (PBS containing 2% FBS and 1 mM EDTA).

Naïve B cells (CD3-CD19+CD27-) were subsequently purified from the PBMC suspension using the EasySep™ Human Naïve B Cell Isolation Kit (Catalog #17254, STEMCELL Technologies) via immunomagnetic negative selection. Following the manufacturer's protocol, the Isolation Cocktail Enhancer (50 µL/mL) and the Naïve B Cell Isolation Cocktail (50 µL/mL) were added to the cell suspension and incubated for 5 minutes at room temperature. EasySep™ Dextran RapidSpheres™ (50 µL/mL) were vortexed and immediately added to the sample. The tube was topped up with recommended medium and placed into an EasySep™ magnet for 3 minutes. The enriched cell suspension was poured off into a new tube. This tube was then placed back into the magnet for a second 1-minute separation. The final supernatant, containing the untouched, highly purified naïve B cells, was collected for subsequent experiments.

#### 4.3 Cell Labeling for Mixing Experiments

To distinguish cell populations in mixing experiments, isolated cells were labeled with fluorescent dyes. Cells were washed and resuspended in pre-warmed, serum-free media (such as PBS) at a concentration of approximately 1.5 to 3 x $10^6$ cells/mL. A stock solution of either CellTracker™ Green CMFDA (Invitrogen, C7025) or CellTracker™ Deep Red (Invitrogen, C34565) dissolved in DMSO was added to the cell suspension at a final working concentration (e.g., 1:1000 for Green, 1:500 for Deep Red). The cell-dye mixture was incubated at 37°C for 20 minutes, with intermittent shaking to ensure uniform labeling. The reaction was quenched by adding an excess volume of complete culture media. Finally, the labeled cells were centrifuged, washed, and resuspended in fresh media at the desired concentration for seeding.

#### 4.4 BLOMB Formation and Stimulation

To induce B cell activation, proliferation, and aggregation into BLOMBs, cells were stimulated with a co-stimulatory cocktail. The primary activation signal was provided by the Human CD40-Ligand Multimer Kit (Miltenyi Biotec, Catalog #130-098-776). Following the kit's protocol, the recombinant CD40-Ligand component was first combined with the multimerization antibody to form an active complex. This complex was then added directly to the cell suspension in complete culture media, along with recombinant cytokines Recombinant Human IL-4 (Gibco, Catalog #CTP0041) and/or Recombinant Human IL-21 (Gibco, Catalog #PHC0211). This combination of CD40 cross-linking and cytokine signaling provides the necessary signals for cellular activation, proliferation, and the subsequent formation of aggregates.

#### 4.5 Image Acquisition and Preprocessing

BLOMBs were imaged at regular intervals (2.5 or 5 minutes per frame) using either an Agilent BioTek Cytation 5 Cell Imaging Multimode Reader or a Zeiss Axio Observer microscope. Both brightfield and fluorescence microscopy modes were used. For mixing experiments, we used differentially labeled cell populations where one population expressed fluorescent markers while the other remained unlabeled. Image stacks were exported as 8-bit TIFF files for subsequent analysis.

#### 4.6 Circularity Measurement from Time-Lapse Data

##### 4.6.1 Automated Segmentation for Population Analysis

For population-level analysis of BLOMB morphology, we employed ilastik [**?**], a machine learning-based image segmentation tool. The pixel classification workflow was trained to distinguish BLOMBs from background using a random forest classifier with features including intensity, edge, and texture filters at multiple scales. The resulting binary masks were processed to extract morphological properties including area, perimeter, and circularity for all detected objects above a minimum size threshold (typically 50 $\mu m^2$).

##### 4.6.2 Semi-Automated Tracking for Individual BLOMB Analysis

To track circularity changes of individual BLOMBs over time, we developed a semi-automated approach. First, the user manually identifies the center of the BLOMB of interest in each frame. A square region centered on this point is then extracted from the full image. Within this local region, Otsu's method is applied to automatically identify the BLOMB boundary. For brightfield images, objects darker than the threshold are selected, while the threshold can be inverted for fluorescent images. If multiple objects are detected in the region, the object closest to the center point is selected as the BLOMB of interest. The circularity metric is then computed as:

$$\text{Circularity} = \frac{4\pi A}{P^2} \tag{10}$$

where $A$ is the object area and $P$ is its perimeter, calculated by counting exposed pixel edges. Finally, the $(x, y)$ coordinates of the detected boundary are stored for each frame, enabling reconstruction of shape evolution. This approach was applied to multiple BLOMBs across different experimental conditions, with typically 10-20 individual BLOMBs tracked per condition.

#### 4.7 Quantification of Cell Mixing at BLOMB Interfaces

To measure the dynamics of cell mixing between differentially labeled populations, we developed a coefficient of variation (CV) based metric that quantifies local heterogeneity in fluorescence intensity.

##### 4.7.1 Interface Region Selection

Circular regions of interest (ROI) were positioned at the interface between fluorescently labeled and unlabeled BLOMBs. To do this, brightfield and fluorescence channels were combined using weighted addition to create grayscale overlay images that clearly show both BLOMB boundaries and fluorescent labeling patterns. For each time point, the user clicks on the interface location where labeled and unlabeled cells meet. A circular region (radius typically 22 pixels, approximately 33 µm) centered on the clicked point is then extracted, and all pixels within this circle are included in the analysis.

##### 4.7.2 Coefficient of Variation Analysis

The mixing metric is computed by first extracting the fluorescence intensities of all pixels within the circular ROI for each time point. The coefficient of variation is then computed as:

$$CV = \frac{\sigma}{\mu} \tag{11}$$

where $\sigma$ is the standard deviation and $\mu$ is the mean of pixel intensities within the ROI.

#### 4.8 Software Implementation

All image analysis routines were implemented in MATLAB R2024a. The analysis pipeline consists of modular scripts for the interactive selection and tracking of regions of interest, automated calculation of morphological and mixing metrics, batch processing of multiple datasets, and statistical analysis. The semi-automated approach balances efficiency with accuracy, enabling analysis of hundreds of BLOMBs while maintaining user oversight. The stored boundary coordinates and ROI positions enable reproducible analysis.
